# Insular Cortex Mediates Approach and Avoidance Responses to Social Affective Stimuli

**DOI:** 10.1101/108506

**Authors:** Morgan M. Rogers-Carter, Juan A. Varela, Katherine B. Gribbons, Anne F. Pierce, Morgan T. McGoey, Maureen Ritchey, John P. Christianson

## Abstract

Social animals detect the affective states of others and utilize this information to orchestrate appropriate social interactions. Social affective behaviors include cooperation, reproductive acts and avoiding sick individuals. In a social affective behavioral test in which experimental adult male rats were given the choice to interact with either naive or stressed conspecifics, the experimental rats demonstrated both approach and avoidant behaviors towards the conspecific, depending upon the age of the conspecific; experimental adult rats approached the stressed juvenile but avoided the stressed adult. Optogenetic inhibition of the insular cortex, a region anatomically positioned to contribute to social cognition, disrupted these behaviors. Receptors for the social nonapeptide oxytocin (OT) are found in high density within the insular cortex and here oxytocin increased intrinsic excitability and synaptic efficacy in acute insular cortex slices. Blockade of oxytocin receptors (OTRs) in the insula eliminated the effect of conspecific stress on approach behavior, while insular administration of OT recapitulated the behaviors typically observed in response to stressed conspecifics. Network analysis using Fos immunoreactivity identified functional connectivity between the insular cortex and the network of regions involved in social decision making. These results implicate insular cortex as a novel target of OT and suggest that insula is a key component in the circuit underlying age-dependent social responses to stressed conspecifics.

---

Social animals, including humans, have an enormous repertoire of behavioral expressions that provide for the transmission of one’s affective state to other members of the group ^1–3^. Sensory and perceptive systems in the “social decision-making network” (SDMN) which consists of the social brain network ^4^ and the mesolimbic reward system ^5^ allow one to appraise these social stimuli and integrate them with past experiences, situational, and somatic factors to shape specific behavioral responses ^6^. In addition to the SDMN, growing evidence implicates insular cortex in responding to socioemotional stimuli in humans. When tasked to identify the emotion of another from a facial expression, or to observe another suffer a painful stimulation, a reliable neural correlate is relative increase in the blood oxygen level dependent (BOLD) signal in the insular cortex ^7,8^. Accordingly, emotion recognition and empathic deficits have been reported among individuals with insular cortex lesions ^9–12^. Empathic processes are also disrupted in autism spectrum disorder (ASD) and fragile X syndrome in which symptoms have been related to insular cortex hypofunction and altered functional connectivity ^13–18^. Translational studies have not yet assessed the role of insula in responding to affective social stimuli and the evidence in support of insula in social cognition from neuroimaging and patient-based studies do not yet demonstrate the necessity of the insula ^19,20^.

The correlation of insular cortex BOLD response and socioemotional cognition in humans is likely a consequence of insular connectivity. The insular cortex has olfactory, gustatory, and somatosensory receptive fields ^21,22^ and is an anatomical locus of sensory integration ^22,23^. There are numerous afferent and efferent tracts connecting the insular cortex to the SDMN, including inputs from the hypothalamus ^24^, ventral tegmental area ^25^, reuniens nucleus ^26^, reciprocal connections with the extended amygdala ^27^ and efferent connections to periaqueductal gray ^28^, bed nucleus of the stria terminalis, nucleus accumbens ^29^ ^30^, basal forebrain ^31^, and medial prefrontal cortex ^32^. The insular cortex also receives input from oxytocin (OT)-containing axons originating from the paraventricular nucleus ^33^ and OT receptor (OTR) binding is particularly enriched in the insular cortex ^34,35^. OT is a critical mediator of reproduction, pair-bonding, and other social behaviors across species ^36–38^ and the presence of OT fibers and OTRs is one of the inclusion criteria for the social brain^4^. Importantly, OT is thought to be an important modulator of empathic cognition in both human ^39,40^ and animal models ^41^.

The OTR is a G-protein coupled receptor which activates the Gα_q11_ signaling cascade ^42^, and OTR activation on insular cortex neurons would provide a mechanism for OT to augment insular cortex activity. Indeed, experimenter-administered OT augments behavioral performance in emotion recognition, relieves some social deficits observed in individuals with ASD; and increases insular activity and connectivity to frontal cortex ^43–45^. Furthermore, mutations in the OTR gene predict ASD symptoms ^46^, insular activation ^47^ and social cognition ^48,49^. In sum, the insular cortex appears critical for social affective behaviors and may be a substrate for pathophysiology in social affective mental illness.

Translational rodent models that capture aspects of social affective cognition include emotion contagion ^50–53^, social buffering ^41^ and even helping ^54,55^ (for review of models see ^56^). However, interpreting the underlying cognitive mechanisms is challenging because some paradigms involve repeated conditioning, direct exposure to a conspecific in pain or receiving aversive stimulation ^57–60^, making it difficult to rule out self-serving or socially rewarding motives ^41,61^. We developed a rat social affective preference (SAP) test in which social affective behaviors were objectively quantified as a preference to approach or avoid interaction with a conspecific that received a mild stressor. Because the experimental rat in the SAP test was not exposed to, nor witness to, the stressor itself, the unconditioned behavior of the experimental subject toward the conspecifics can be interpreted as a response to the affective state of the target. Using the SAP test, we report a set of *in vivo* and *in vitro* studies that tested whether insular cortex activity and modulation by OT were necessary and sufficient to modulate social affective behaviors in rat. The results clearly implicate the insula in responding to stressed conspecifics. Finally, we employed graph-theory based network analyses ^62^ to characterize, in rats, the relationship of the insular cortex to several regions of interest (ROIs) implicated in social decision making.

## RESULTS

### Age and stress exposure of conspecific determine social approach

In the social affective preference (SAP) test an experimentally naive, adult male rat was presented with a pair of unfamiliar conspecific male stimuli (see Fig. 1 and Supplementary Video 1). To manipulate social affect, one of the conspecifics was exposed to a mild stressor of 2, 5s 1mA footshocks immediately before the test, while the other did not receive footshocks (naïve). Because social approach behaviors are shaped, in part, by features of the target including age ^63^, we hypothesized that rats may differentially respond to stressed conspecifics as a function of the target’s age. Experimental adult male rats (PN 60-80 days) underwent SAP tests (See Figs. 1A & 1B) in which the choice test involved exposure to a pair (one stressed, one naive) of unfamiliar pre-pubertal male juveniles or a pair of unfamiliar male post-pubertal adults ^64^. The design was a 2 by 2 with conspecific Age (PN 30 vs. PN 50) as a between-subjects factor and conspecific Affect (naive or stress) as a within-subjects factor. We observed a significant increase in social interaction with the stressed juvenile (*p* < 0.001) but a significant decrease in interaction with the stressed adult (*p* < 0.05, Fig. 1C). To summarize this interaction, we computed a percent preference score by dividing the time spent interacting with the stressed conspecific by the total time spent interacting during the test (Fig. 1D). Thus, when faced with a choice to interact with either a naive or stressed conspecific, we observed a preference to interact with a stressed juvenile but avoidance of a stressed adult. Analysis of *conspecific* behavior (Supp. Fig. 1) revealed that stress caused juvenile conspecifics to engage in more self-grooming compared to the naive juvenile conspecific, but no other behaviors differed significantly between stressed and naïve rats in either the juvenile or adult groups.

**Figure 1.**
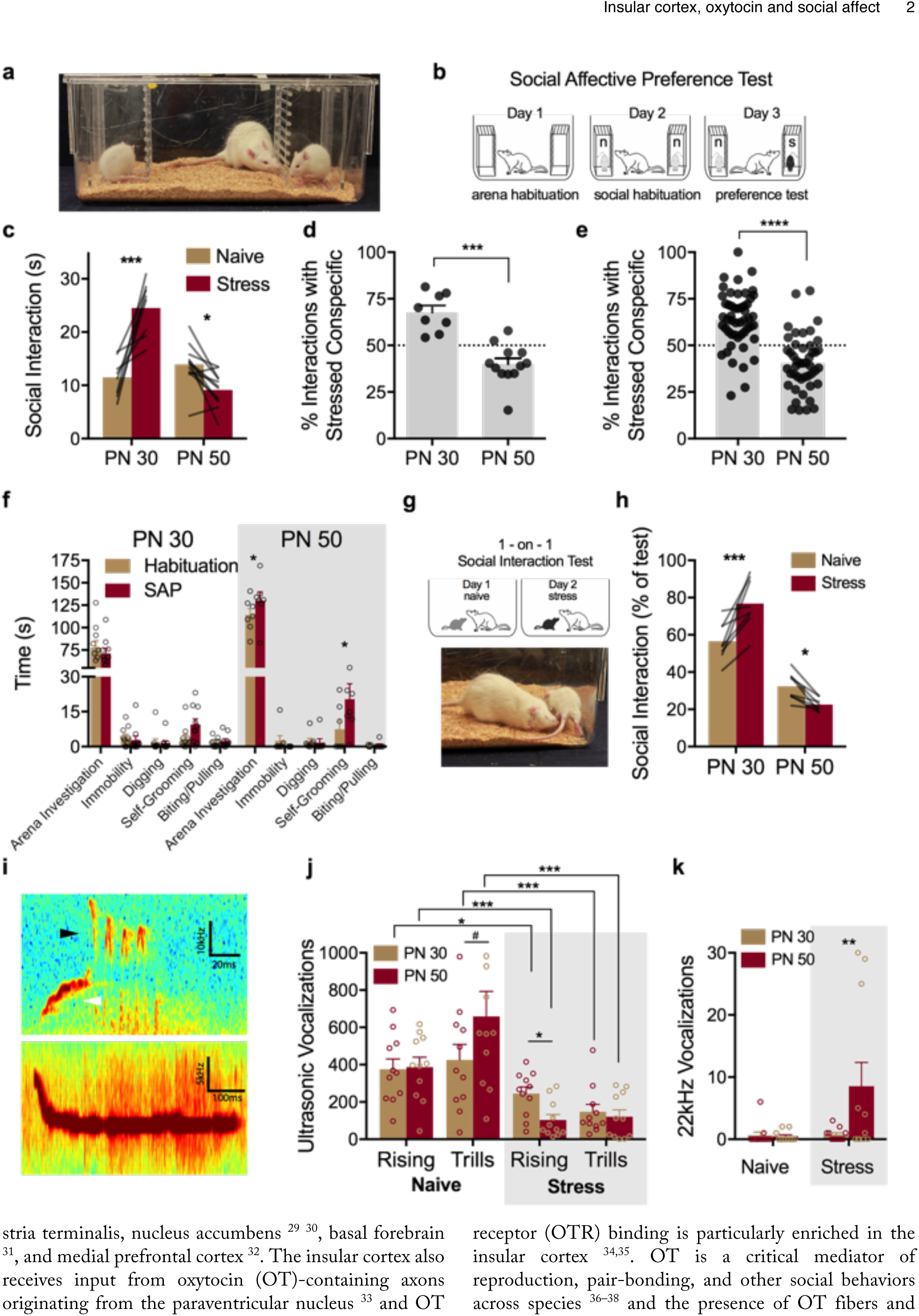
Social affective preference (SAP) **(A)** The SAP arena containing juvenile conspecifics on the left and right with an experimental adult in the center. **(B)** Diagram of SAP test procedure. Experimental adults were allowed to explore the test arena and conspecific chambers when empty (Day 1), with two experimentally naive conspecifics (Day 2) and when given a choice to interact with either a naive (n) or recently shocked (s) conspecific (Day 3). **(C)** Mean (individual replicates shown as connecting lines) time spent in social interaction with the naive or stressed conspecific by age (n = 8, PN 30; n = 12, PN 50). Although conspecific age did not alter time spent interacting with the naive conspecific, a bidirectional effect of age was apparent in time spent interacting with the stressed conspecifics (*F*_AGE_(1, 18) = 27.93, *p* < 0.001; *F*_AFFECT_(1, 18) = 9.965, *p* = 0.006; *F*_AGE*AFFECT_(1, 18) = 46.05, *p* < 0.001). Experimental rats spent more time exploring the stressed PN 30 conspecific compared to the PN 30 naive, but spent less time exploring the stressed PN 50 conspecific compared to the PN 50 naive. For analyses of conspecific behavior please refer to Supplementary Figure 1. **(D)** Mean (+ S.E.M. with individual replicates) data in C expressed as the percent of time spent interacting with the stressed conspecifics relative to the total time spent interacting. Here, experimental adults showed a marked preference (values greater than 50%) for interaction with stressed conspecifics and avoidance (values less than 50%) of stressed adults (t(18) = 5.783, *p* < 0.001). **(E)** Mean (+ S.E.M. with individual replicates, *n* = 51 for PN 30, *n* = 46 for PN 50) percent preference for interacting with the stressed conspecific pooled from all of the subjects in the experimental control groups included in the current report including vehicle, sham, light OFF control groups of the later experiments. Percent preference scores for PN 30 and PN 50 interactions were significantly different from each other (*t*(95) = 7.66, *p* < 0.0001), and in both conditions the mean percent preference differed from 50% (PN 30: *t*_one-sample_(50) = 6.49, *p* < 0.0001; PN 50: *t*_one-_sample (45) = 4.39, *p* < 0.0001). **(F)** Mean (+ S.E.M. with individual replicates) time spent in non-social behaviors during habituation tests and SAP tests (n = 10, PN 30; n = 7 PN 50). More investigation of the arena and time spent self-grooming was observed in the PN 50 rats (*F*_AGE*BEHAVIOR*TEST_(4, 60) = 3.014, *p* < 0.025). **(G)** Diagram of 1-on-1 social interaction test and photo of typical adult-initiated interactions. Experimental adult rats were presented on Day 1 with a naive conspecific and experimental adult-initiated interactions were recorded for 5 min. On the next day, the conspecific received 2 footshocks immediately prior to the test; order of testing was counterbalanced. **(H)** Mean (individual replicates, *n*s = 8/ age) time interacting with the naive or stressed conspecific in a 1-on-1 test shown as percent of test time. Experimental adults spent significantly more time interacting with the stressed PN 30 conspecific but significantly less time with the stressed PN 50 conspecific compared to the respective naive conspecific targets (*F*_AGE_(1, 14) 103.10, *p* < 0.001; *F*_AGE*AFFECT_(1, 14) = 31.34, *p* < 0.001). **(I)** Representative audio spectrograms depicting rising (top, white arrow), trills (top, black arrow) and 22kHz ultrasonic vocalizations (USVs). Scale bars indicate Y-axis ranges: 60-70kHz (top) and 30-35kHz (bottom). **(J-K)** Mean (+S.E.M with individual replicates, *n*s = 11/group) number of rising and trill USVs recorded during 5 min one-on-one social interactions. Fewer rising and trill calls were observed during interactions with stressed conspecifics with fewer rising calls observed in stressed adults compared to stressed juveniles but more 22kHz calls observed in stressed adults than stressed juveniles (*F*_STRESS_(1, 40) = 26.16, *p* < 0.001; *F*_CALL TYPE_(2, 80) = 60.86, *p* < 0.001; *F*_STRESS*CALL TYPE_(2, 80) = 20.18, *p* < 0.001; *F*_CALL TYPE*AGE_(2, 80) = 3.43, *p* = 0.37). \**p* < 0.05, ** *p* < 0.0, *** *p* < 0.001, **** *p* < 0.0001

The SAP procedure was replicated several times in the later optogenetic and pharmacology experiments which provided a large sample to evaluate the reliability and generality of these phenomena, and to quantify changes in experimental adult rat behavior from the habituation day to the SAP test day (videos of both were not recorded in first experiments). Data from SAP tests conducted under either vehicle, Light-OFF or sham treatments were pooled and converted to percent preference scores (Fig. 1E) and the scores were found to fit a normal distribution (D’Agostino and Pearson normality test, for PN 30: *K2* = 2.52, *p* = 0.28, for PN 50: *K2* = 2.14, *p* = 0.34). One sample t-tests were used to compare PN 30 and PN 50 preference scores to the hypothetical value of 50% (equal time exploring both naive and stressed conspecifics) revealed a preference for the stressed PN 30 (*p* < 0.0001) and a preference for the naive PN 50 (*p* < 0.0001). Approximately 82% (42 of 51) of rats tested with PN 30 conspecifics exhibited a preference for the stressed target, while only 21% (10 of 46) of rats tested with PN 50 conspecifics exhibited preference for the stressed target. To evaluate whether exposure to the stressed conspecific in the SAP tests altered any aspect of experimental adult behavior, we quantified behaviors (detailed in Online Methods) in the experimental adult rats during habituation and SAP tests with either PN 30 or PN 50 conspecifics (Fig. 1F). Videos were selected blind to SAP test social interaction results. A significant Age by Behavior by Test (habituation vs. SAP) interaction (*p* = 0.025) reflected increases in arena investigation (*p* = 0.05) and self-grooming (*p* = 0.035) during SAP tests with PN 50 conspecifics.

The SAP results predict that in a one-on-one interaction an experimental adult should engage in more social interaction when exposed to a stressed juvenile than when exposed to a naïve juvenile, and the opposite pattern for adult targets. Thus, experimental adult rats were given a series of 2 “one-on-one” social exploration tests (5 min duration, one test per day) with either 1 unfamiliar naive PN 30 juvenile, or 1 unfamiliar stressed juvenile as the social interaction target (Fig. 1G); a separate set of experimental adults received the same series of tests with PN 50 adult conspecifics. A within-subjects design was used with test order counter balanced and stress treatment was as above. As in the SAP test, experimental adults spent more time interacting with the stressed juvenile (*p* < 0.001) and less time with the stressed adult (*p* = 0.043; Fig. 1H). Because this effect could be influenced by differential conspecific-initiated social interaction, i.e. reciprocal interactions, we also quantified exploration of the experimental adult by the conspecific; however, there was no difference between naive-to-adult or stressed-to-adult interactions at either age.

#### Conspecific stress alters ultrasonic vocalizations

Ultrasonic vocalizations (USVs) have been reported to signal emotional states in rats ^65^. To explore whether USVs differed during social interactions with stressed conspecifics, a separate set of rats was given a single, 5 min one-on-one social interaction test in a chamber equipped with a lid-mounted ultrasonic microphone and social interaction time and USVs were quantified during exposure to one of 4 conspecific stimuli: Naive PN 30, Naive PN 50, Stressed PN 30 or Stressed PN 50. Three types of vocalizations were present in our recordings: 22kHz, flat vocalizations, ~30-60kHz rising calls, and ~60-80kHz trills (Fig. 1I). 22kHz vocalizations are thought to convey negative affect, and these were observed primarily during interactions with stressed adults. Rising and Trill calls, which are emitted during and in anticipation of rewarding stimuli, were quite frequent during naive interactions but reduced during stressed interactions. Post hoc comparisons revealed more 22kHz calls in the PN 50 stress rats than PN 30 stress rats or PN50 naive rats (*p*s < 0.007); fewer rising calls in PN 30 and PN 50 stress rats compared to naive levels (*p*s < 0.045), more rising calls in the stressed PN 30 than stressed PN 50 (*p* = 0.03), fewer trills in the PN 30 and PN 50 stress conditions than naive (*p*s < 0.02) and a trend for more trills in the naive PN 50 compared to PN 30 (*p* = 0.055). Although it is impossible to determine whether stress altered USVs emitted by conspecifics, USVs emitted by the experimental rat toward the stressed conspecific, or some combination of both, stressor exposure dramatically shifted the patterns of USVs consistent with a state of negative affect.

#### Optogenetic silencing of insular cortex prevented social affective preference

To test whether exposure to stressed conspecifics influences insular cortex activity, experimental rats were sacrificed 90 min after the one-on-one social interaction tests described above for USV analysis and sections containing the insular cortex were stained for Fos immunoreactivity (Fig. 2A). Fos immunoreactivity was higher in the insula of rats that had interacted with stressed PN 30 conspecifics than in rats that had interacted with stressed PN 50 conspecifics (*p* = 0.009), and insula Fos immunoreactivity was lower in rats after interaction with PN 50 stressed rats than in rats after interaction with PN 50 naive rats (*p* = 0.038; pooled across region). To determine how insular cortex Fos counts relate to social interaction we conducted a linear regression analysis with Age and Stress conditions included as moderators of insula Fos levels (Fig. 2C). The results indicate large effects of insula Fos levels (*p* < 0.001), Age (*p* = 0.001) and marginally significant interactions of insula Fos levels by Stress (*p* = 0.055) Stress by Age (*p* = 0.051). Insular cortex activity, as indexed by Fos immunoreactivity, reflects both the age and stress state of the conspecific. To test whether insula activity is necessary for social affective behaviors we conducted a pilot study in which pharmacological inhibition of the insula prior to SAP tests altered the pattern of behavior observed in experimental rats (Supplementary Fig. 3). This suggests that the activity and, perhaps, output of the insular cortex is necessary for normal social affective behavior.

**Figure 2.**
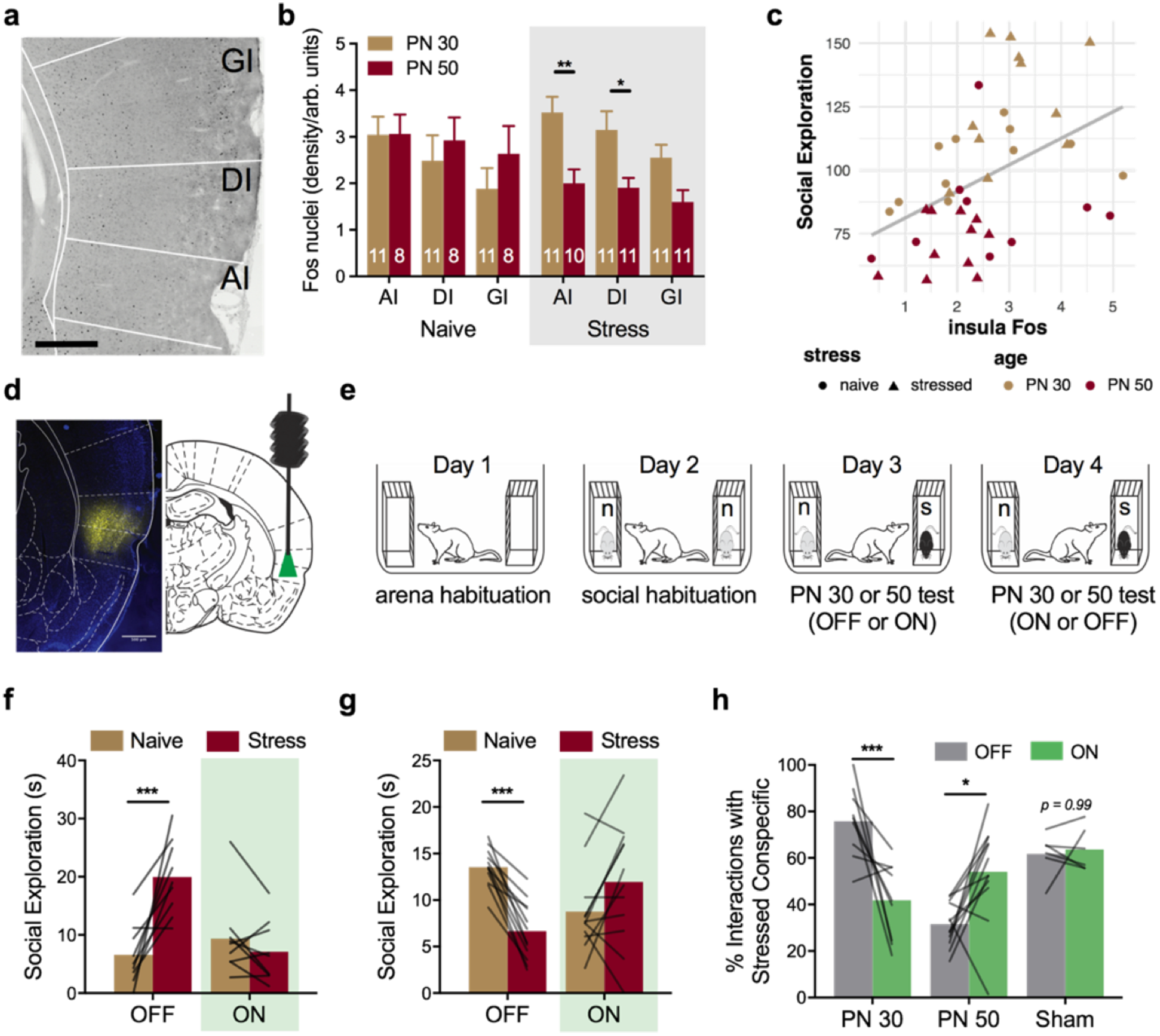
Optogenetic silencing of insular cortex during SAP tests. **(A)** Representative digital photomicrograph containing insular cortex regions and Fos immunoreactive nuclei (black ovoid particles). Scale bar = 500µm. **(B)** Mean (+S.E.M.) Fos immunoreactive nuclei by insular cortex subregion (AI = Agranular, DI = Dysgranular, GI = Granular) quantified 90 min after social interaction with a naive PN 30, naive PN 50, stressed PN 30 or stressed PN 50 conspecific (5 min test). Fos was found in all regions but there was an effect of stress in the PN 50 groups such that less Fos was evident in the PN 50 brains after stressed conspecific interactions than in PN 50 brains after naive conspecific interactions, and there was less Fos in the AI and DI in the PN 50 Stressed group compared to the PN 30 stressed group (*F*_STRESS*AGE_(1, 36) = 6.21, *p* = 0.017; *F*_SUBREGION_(2, 72) = 7.90, *p* = 0.001). **(C)** Mean Fos immunoreactivity (pooled across insula) predicted time spent in social interaction during the 5 minute test. Regression analysis indicated strong prediction of social interaction by Fos level, Age, and interactions between Stress, Fos and Age (*F*_FOS_(1, 34) = 19.72, *p* < 0.001, *F*_AGE_(1, 34) = 36.93, *p* < 0.001; *F*_FOS*STRESS_(1, 34) = 3.97, *p* = 0.055; *F*_STRESS*AGE_(1, 34) = 4.09, *p* = 0.051). To test whether the insular cortex is required for social affective behavior, rats were transduced with a viral construct containing eNpHr3.0 fused to mCherry in the insular cortex and implanted with bilateral fiber optic cannula guides terminating at the dorsal border of the insular cortex. **(D)** Native (unamplified) mCherry expression in the insular cortex (false colored yellow) from a brain slice adjacent to one containing the cannula tract imaged at 10x. (Scale bar = 500um). **(E)** Diagram of SAP tests for optogenetic experiments. On Day 1 rats were habituated to the arena. On Day 2 rats were handled to habituate to connecting fibers and exposed to the arena with naive conspecific stimuli. On Days 3 and 4, optical fibers were inserted bilaterally and SAP tests were performed under no light (OFF) or continuous green light (ON) conditions; order was counterbalanced. **(F)** Mean (individual replicates, *n* = 9) time spent interacting with PN 30 juvenile conspecifics on Days 3 and 4 of the SAP test. In the light OFF condition, the experimental adult spent significantly more time interacting with the stressed conspecific but this pattern was abolished in the light ON condition (*F*_AGE*STRESS*LIGHT_(1, 19) = 41.31, *p* < 0.001). **(G)** Mean (individual replicates, n = 12) time spent interacting with PN 50 adult conspecifics on Days 3 and 4 of the SAP test. In the light OFF condition, the experimental adult spent significantly less time interacting with the stressed conspecific but this pattern was reversed in the light ON condition. **(H)** Data from F and G converted to percent preference for interaction with stressed conspecifics for comparison with a sham control group that did not express eNpHr3.0 and was later given SAP tests with PN 30 conspecifics. Here a clear age by light interaction is apparent with optogenetic silencing of insular cortex eliminating preference for interaction with the stressed juvenile, and blocking the pattern of avoidance of stressed adult conspecifics. Light had no effect in the shams. * *p* < 0.05, \*\**p* < 0.01, *** *p* < 0.001.

To achieve inhibition of insular cortex pyramidal neurons, rats were transduced with halorhodopsin (eNpHr3.0) under the CamKII promoter (AAV5-CamKIIα-eNpHR3.0-mCherry), which allowed reversible neuronal silencing (Fig. 2D) ^66^. Optogenetic transductions were evaluated by fluorescence microscopy and *in vitro* whole cell recordings (Supplementary Fig. 4). Two weeks after virus infusion, rats underwent SAP tests in a 2 Age (juvenile or adult conspecific) by 2 Affect (naive or stress) by 2 Light (OFF or ON) design (Fig. 2E). Green light (ON) or no light (OFF) was delivered to the insula continuously during the SAP test. As expected, the majority of rats in the OFF condition exhibited preference to interact with the stressed juvenile (9 of 12 experimental rats) and avoided the stressed adult (12 of 14 experimental rats), whereas experimental rats in the light ON condition avoided the stressed juvenile (Fig. 2F) or preferred interaction with the stressed adult (Fig. 2G) (Age*Stress*Light interaction, *p* < 0.001). The 5 rats that did not exhibit the expected preference pattern in light OFF were analyzed separately (Supplementary Fig. 5). Pairwise comparisons in the juvenile OFF condition identified an increase in interaction of the stressed juvenile compared to naive juvenile (*p* < 0.001), whereas in the adult OFF condition there was a significant decrease in interaction with the stressed adult conspecific compared to the naive adult conspecific (*p* = 0.01). In the light ON condition, there were no significant differences in exploration times between naive and stressed conspecifics in either age condition (PN 30 *p* = 0.22 and PN 50 *p* = 0.054, approaching significance for the opposite direction of the light OFF condition). This indicates that optogenetic silencing of the insular cortex prevented the expression of social affective behaviors. Importantly, optical treatment had no effect on rats with sham transfections when interacting with PN 30 conspecifics (Fig. 2H), and light condition did not influence exploratory activity or general behavior in the SAP test (Sup. Fig. 4).

#### Oxytocin alters the excitability of the insular cortex

The foregoing suggested that exposure to stressed conspecifics triggers a neurobiological response that alters insular cortex activity. Given the dense expression of OTR in the insula and evidence that OT can modulate excitatory balance in cortical circuits ^67^, we hypothesized that OT released during the SAP test could directly modulate insular cortex excitability. The excitability of insular cortex was characterized after bath application of OT (500nM) in whole-cell pyramidal neuron recordings (Fig. 3) and in extracellular multiple electrode field excitatory postsynaptic potentials (fEPSPs) in acute slices (Fig. 4). Active and passive intrinsic properties were quantified, as previously ^68^. A complete list of parameters measured and results of paired samples t-tests is provided in Table 1.

**Figure 3.**
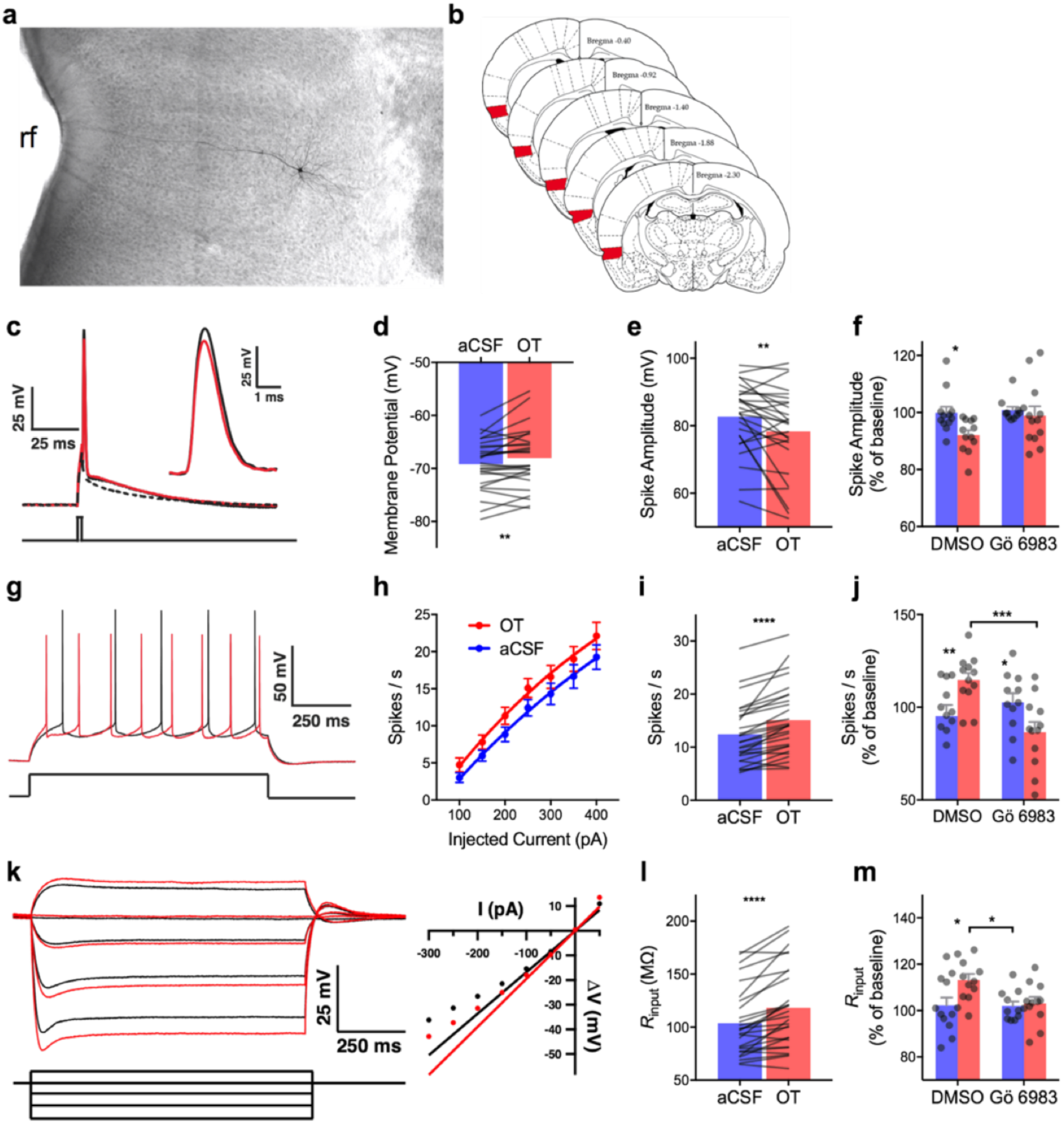
Intrinsic membrane properties in insular cortex pyramidal neurons are modulated by oxytocin via protein kinase C (PKC). **(A)** Digital photomicrograph of a typical biocytin filled neuron, rf = rhinal fissure. **(B)** Schematic diagram illustrating (red shading) the region of interest for whole cell recordings. All neurons included in these experiments were found in this region. **(C)** Typical action potential before (black trace) and after bath application of 500nM OT (red). Inset: detail of the AP peak amplitude difference. Intrinsic properties were characterized from a sample of 27 neurons; the dependence of OT effects on PKC was determined by first characterizing the change in intrinsic properties from baseline followed by either no treatment (aCSF) or OT (500nM) in either the presence of DMSO or DMSO and the pan-PKC inhibitor Gö 6983 (200nM, *n* = 11 to 12 neurons/group). **(D)** Mean action potential amplitude; OT significantly reduced amplitude. **(E)** Mean resting membrane potential. OT significantly depolarized the membrane at rest. **(F)** Mean action potential amplitude (+S.E.M. with individual replicates); OT significantly reduced spike amplitude but did not change in the presence of Gö 6983 (*F*_OT_(1, 21) = 4.56, *p* = 0.044, aCSF vs. OT, *p* = 0.040). **(G)** Typical train of spikes evoked by 1 s 150pA current injection before (black) and after (red) OT. **(H)** Mean (+/− SEM) spikes evoked by increasing current injections; OT increased the spike frequency (*F*_OT_(1, 279) = 10.42, *p* = 0.001). **(I)** Mean action potentials evoked by 250pA current; significantly more spikes were evoked after OT. **(J)** Mean (+ S.E.M. and individual replicates) spike frequency upon 250pA depolarization; OT increased spiking compared to aCSF and Gö 6983 reduced spiking on its own (*F*_OT*Go_(1, 21) = 16.44, *p* < 0.001, aCSF-OT vs. Gö-OT, p < 0.001). **(K)** Membrane potentials evoked by subthreshold and hyperpolarizing current injections (left) and typical rectification curve (right) before (black traces) and after (red traces) OT. **(L)** Mean input resistance (*R*_input_); OT significantly increased *R*_input_. **(M)** Mean (+ S.E.M. and individual replicates) input resistance. OT increased input resistance (*F*_Go_(2, 20) = 4.66, *p* = 0.043, OT-DMSO vs. OT-Gö, *p* = 0.03). Symbols and connecting lines indicate individual replicates. \**p* < 0.05, ** *p* < 0.01, **** *p* < 0.0001.

**Table 1.**
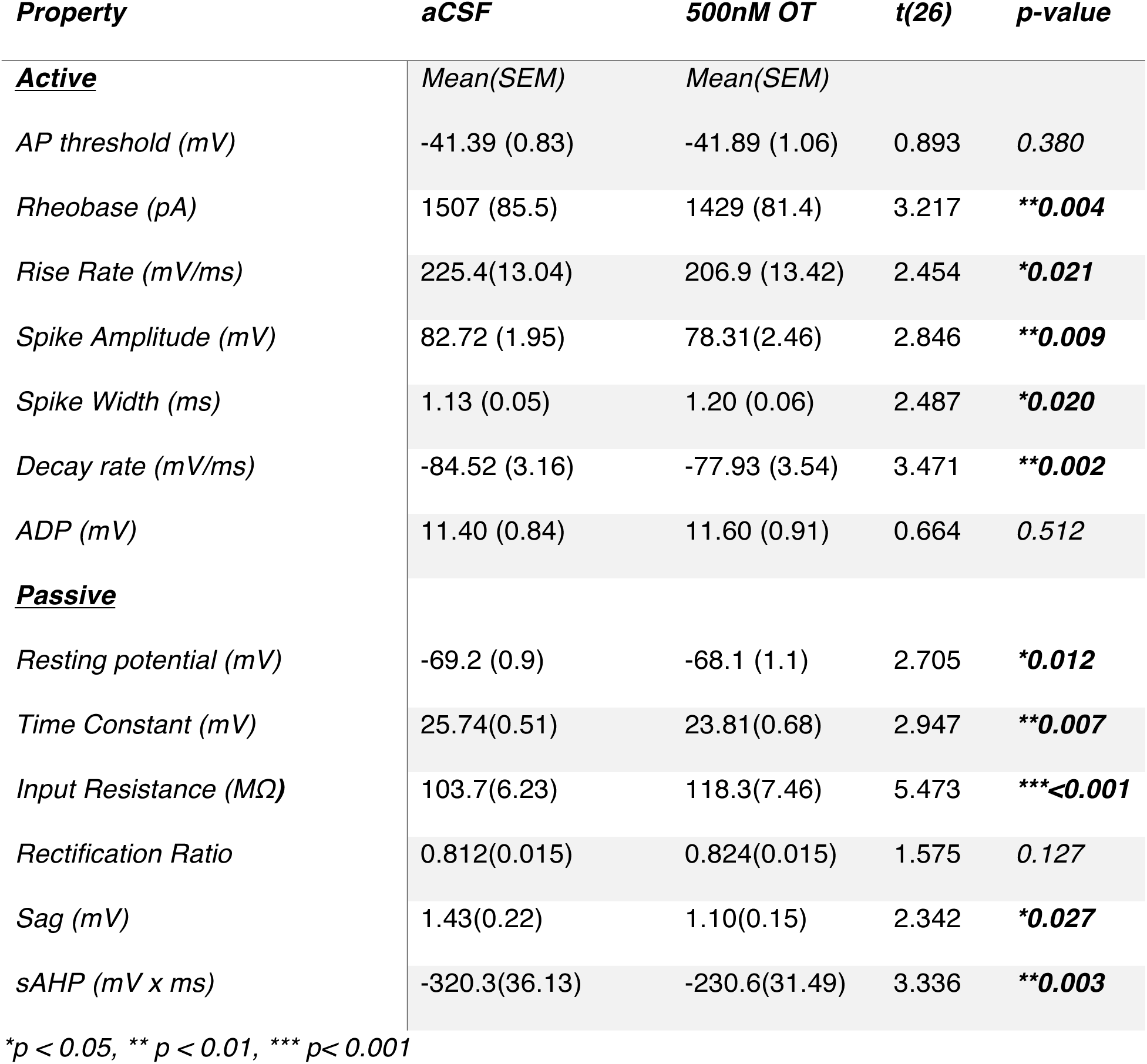
Effects of 500nM OT on insular cortex pyramidal intrinsic membrane properties

| *Property* | *aCSF* | *500nM OT* | *t(26)* | *p-value* |
| --- | --- | --- | --- | --- |
| **<u>Active</u>** | *Mean(SEM)* | *Mean(SEM)* |  |  |
| *AP threshold (mV)* | -41.39 (0.83) | -41.89 (1.06) | 0.893 | *0.380* |
| *Rheobase (pA)* | 1507 (85.5) | 1429 (81.4) | 3.217 | **\*\*0.004** |
| *Rise Rate (mV/ms)* | 225.4(13.04) | 206.9 (13.42) | 2.454 | **\*0.021** |
| *Spike Amplitude (mV)* | 82.72 (1.95) | 78.31(2.46) | 2.846 | **\*\*0.009** |
| *Spike Width (ms)* | 1.13 (0.05) | 1.20 (0.06) | 2.487 | **\*0.020** |
| *Decay rate (mV/ms)* | -84.52 (3.16) | -77.93 (3.54) | 3.471 | **\*\*0.002** |
| *ADP (mV)* | 11.40 (0.84) | 11.60 (0.91) | 0.664 | *0.512* |
| **<u>Passive</u>** |  |  |  |  |
| *Resting potential (mV)* | -69.2 (0.9) | -68.1 (1.1) | 2.705 | **\*0.012** |
| *Time Constant (mV)* | 25.74(0.51) | 23.81(0.68) | 2.947 | **\*\*0.007** |
| *Input Resistance (MΩ)* | 103.7(6.23) | 118.3(7.46) | 5.473 | **\*\*\*<0.001** |
| *Rectification Ratio* | 0.812(0.015) | 0.824(0.015) | 1.575 | *0.127* |
| *Sag (mV)* | 1.43(0.22) | 1.10(0.15) | 2.342 | **\*0.027** |
| *sAHP (mV x ms)* | -320.3(36.13) | -230.6(31.49) | 3.336 | **\*\*0.003** |

\* $p < 0.05$ , \*\* $p < 0.01$ , \*\*\* $p < 0.001$

Figures 3A & 3B depict a typical insular cortex pyramidal neuron and the rostrocaudal extent of insular cortex targeted in this study. All of the analyzed neurons (N = 27) were localized in the deep layers of the agranular insular cortex, where OTR binding is dense ^34,35^. A number of intrinsic properties were altered by application of OT, which together suggested an increase in excitability. Specifically, we observed a depolarization of the resting membrane potential, an increase in input resistance, and a decrease in membrane time constant. These changes would permit the membrane to charge faster and reach the action potential threshold with less excitatory input. Indeed, OT also reduced the minimum current injection required to elicit an action potential (*i.e.* the rheobase). We also observed a significant reduction in the slow after-hyperpolarization (sAHP), quantified as the area (mV x ms) of the hyperpolarization (relative to pre-step baseline) immediately following trains containing equal number of action potentials. Together these changes suggest that after OT release, insular cortex pyramidal cells might achieve greater spike frequency. Consistent with this prediction, spike frequency increased in response to sustained depolarizing currents (Fig. 3G-I). Interestingly, after OT application, action potentials were significantly smaller (Fig. 3C-E) with decreased amplitude, rise rate, decay rate and longer duration (half-width), which may indicate that the increase in firing frequency could come at a cost of reduced neurotransmitter release. OTR is thought to signal through the Gα_q11_ signaling cascade to raise intracellular Ca^2+^ levels and activate protein kinase c (PKC) ^42^. To test whether the changes in intrinsic and synaptic physiology caused by OT are mediated by PKC we applied the pan-PKC antagonist Gö 6983 (200nM) ^69^ in whole-cell recordings with or without OT (500nM). Gö 6983 prevented the decrease in spike amplitude (Fig. 3F), increase in firing frequency (Fig. 3J) and increase in *R*_input_ (Fig. 3M) that were evident in OT treated neurons.

We next investigated the effect of OT application on evoked synaptic efficacy. Input/output curves were conducted in acute insular cortex slices on a perforated 60 channel multiple electrode array (Fig. 4A). fEPSPs were generated by applying biphasic voltage steps to one of the extracellular electrodes located within the deep layers of insular cortex. OT application induced a leftward shift, with significantly larger fEPSP amplitude compared to baseline at stimuli from 2 to 5V. This effect increased during the washout period (Fig. 4B). Comparisons between conditions at each stimulus intensity revealed significant increases in fEPSP amplitude during OT and washout compared to baseline beginning at 2V, with washout levels differing from OT beginning at 3V (*p*s < 0.001). In the aCSF control slices synaptic responses were stable across all phases of the experiment. As above, in the presence of Gö 6983 (200nM), OT (1uM) did not alter fEPSP input/output curves (Fig. 4G-H).

**Figure 4.**
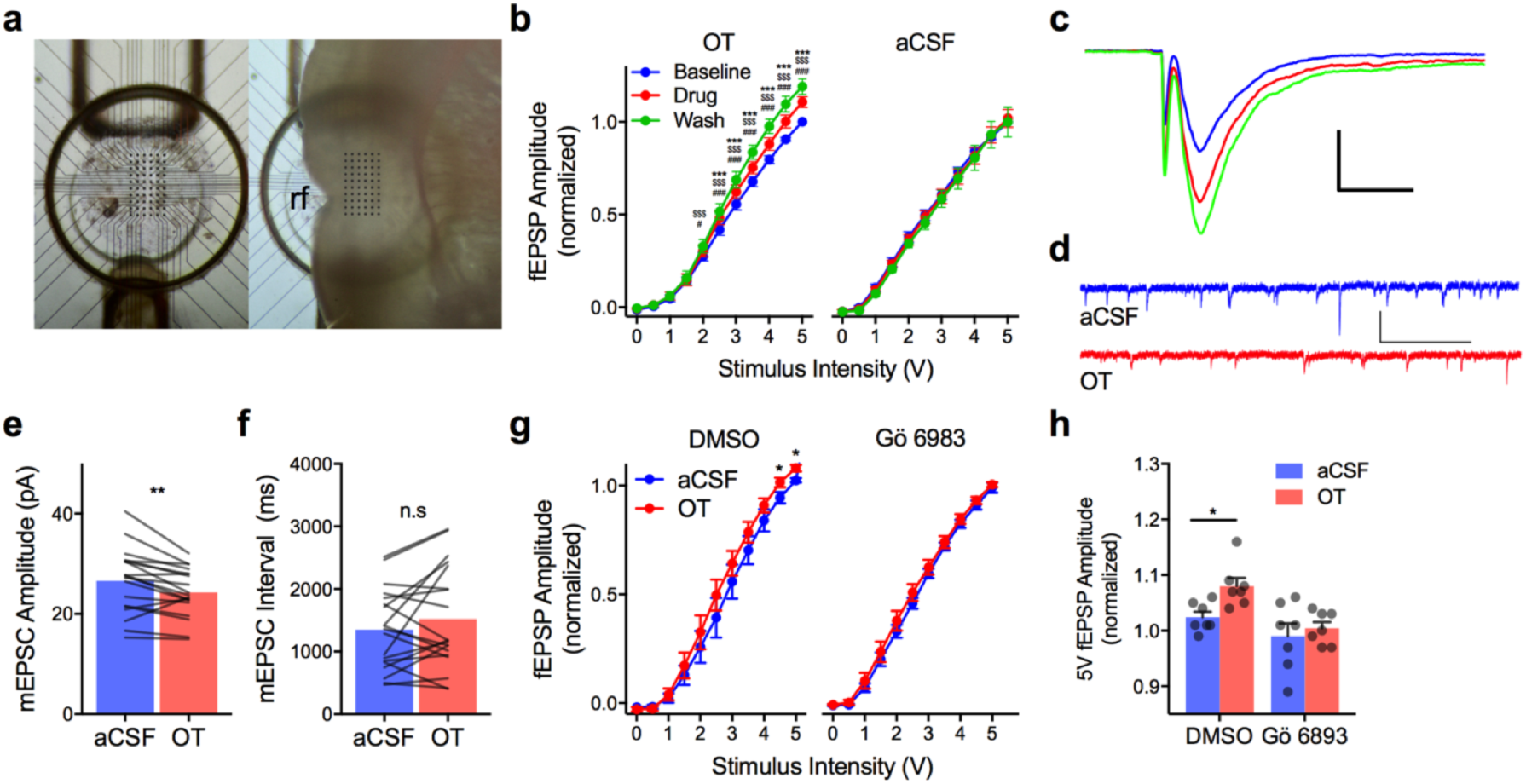
Oxytocin modulates excitatory synaptic transmission in the insular cortex. **(A)** Top view of 60 channel perforated MEA (left) for acute extracellular recordings of the insular cortex. Right depicts location of insular cortex slice during recording, rf = rhinal fissure. **(B)** Input/output curves for fEPSPs (Mean +/− SEM, OT n = 10, aCSF n = 9 slices) normalized to the peak amplitude observed in response to 5V stimulation under baseline conditions. OT significantly increased EPSP amplitude beginning at 2V with further enhancement during the washout (*F*_STIMULUS_(10, 90) = 598.20, *p* < 0.0001; *F*_DRUG_(2, 18) = 11.99, *p* < 0.001; *F*_STIMULUS*DRUG_(20, 180) = 11.34, *p* < 0.0001). Without application of OT, EPSPs remain stable across the duration of the experiment (aCSF: *F*_STIMULUS_(10, 80) = 385.90, *p* < 0.0001). # *p* < 0.05 OT vs. Baseline, ### *p* < 0.001 OT vs. Baseline, $$$ *p* < 0.001 Wash vs. Baseline, *** *p* < 0.001 Wash vs. OT. **(C)** Typical fEPSPs evoked by biphasic extracellular stimulation at baseline (blue) during application of 500nM OT (red) and after washout (green). Scale bar 500μV/ms. **(D)** Representative voltage clamp recordings of mEPSCs recorded before (aCSF; blue) and after OT (red). **(E)** Mean mEPSC amplitude before and after OT (*n* = 19 neurons); OT significantly reduced amplitude, **(paired *t*(18) = 3.29, *p* = 0.004). **(F)** Mean mEPSC interval (n = 19 neurons); no effect of OT was apparent, (paired t(18) = 1.42, *p* = 0.17). **(G)** Input/output curve for fEPSPs (Mean +/− S.E.M., *n* = 7 slices/condition) normalized to the peak amplitude observed in response to 5V stimulation under baseline conditions. OT (1uM) increased fEPSP at 4.5 and 5mV while no effect was observed in the presence of Gö 6983. \**F*_STIMULUS*DRUG_(10, 240) = 3.40, *p* < 0.001, OT vs. aCSF at 4.5 and 5V, *p*s < 0.031. **(H)** Mean (+ S.E.M. and individual replicates) fEPSP amplitude from (G) at 5V.

An increase in intrinsic excitability might also be reflected in a change in the miniature and/or spontaneous release of excitatory neurotransmitter in the slice. We made continuous whole-cell, voltage-clamp recordings of deep layer insular cortex pyramidal neurons with either tetrodotoxin (*n* = 7) or without (*n* = 12) before and after application of OT (500nM) to quantify miniature and spontaneous excitatory postsynaptic currents (EPSCs), respectively. However, in our recording conditions very few large events, indicative of spontaneous glutamate release, occurred, so all events were assumed to be miniature EPSCs (mEPSCs) and the data were pooled (*N* = 19). OT reduced mEPSC amplitude (*p* < 0.01) but had no effect on mEPSC frequency. In a pilot experiment, no effect of OT was found on spontaneous inhibitory post-synaptic potentials (data not shown). A change in mEPSC amplitude likely reflects a postsynaptic modulation of AMPA receptor mediated currents ^70^ but not an effect of OT on the spontaneous circuit activity *per se.* However, no effect of OT was observed in AMPA:NMDA current ratio (data not shown). Taken together, the electrophysiology suggests that OT rendered insular cortex pyramidal neurons more responsive to excitatory inputs via activation of PKC second messenger cascades.

#### Insular cortex oxytocin receptors are necessary for both prosocial and asocial responses to stressed conspecifics

We next determined whether insular cortex OTRs contribute to the prosocial or asocial behaviors in the SAP test. To this end, 20 experimental rats were implanted with cannula guides to the insular cortex. After recovery, the rats underwent the SAP procedure, with preference tests on days 3 and 4 in a repeated measures design (Fig. 5A). Here, the experimental design was a 2 by 2 by 2 with conspecific Age (PN 30 vs. PN 50) as a between-subjects factor, and with Drug (Vehicle or OTR antagonist (OTRa) 20ng/side as in ^71^) and conspecific Stress as within-subjects factors. The rats received microinjections 15 min before SAP tests on days 3 and 4, with drug order counter balanced. The conspecific stimuli were always unfamiliar; no effect of test order was apparent. Two subjects in the PN 50 condition with misplaced cannula were excluded (for cannula placement see Supplementary Fig. 6).

**Figure 5.**
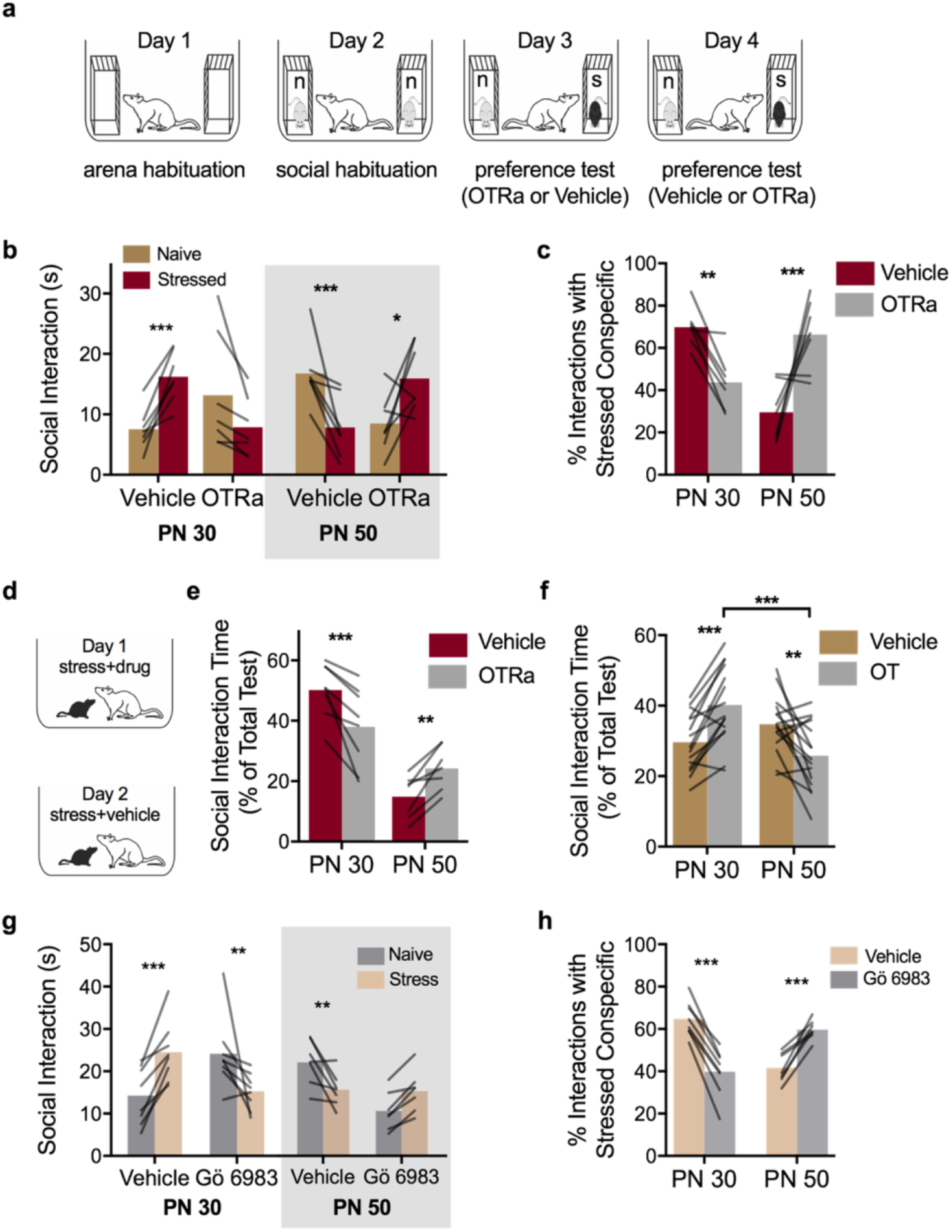
Social affective behaviors require insular cortex oxytocin. **(A)** Diagram of experimental design. **(B)** Mean (individual replicates) time spent exploring either a naive conspecific or a stressed conspecific in the SAP test with either PN 30 (*n* = 7) or PN 50 (*n* = 7) conspecifics after intra-insula infusion of a selective OTR antagonist (OTRa, 20ng/side). Vehicle treated experimental adult rats spent more time interacting with the stressed PN 30 juvenile conspecifics and less time with the stressed PN 50 adult conspecifics. These trends were blocked and reversed by infusion of OTRa (*F*_AGE*DRUG*STRESS_(1, 12) = 31.84, *p* < 0.001). **(C)** Data in (B) expressed as percent preference for interaction with the stressed conspecific (Mean individual replicates). OTRa significantly reduced preference for the stressed PN 30 while increasing time spent with the stressed PN 50 conspecific (*F*_AGE*DRUG_(1, 12) = 26.38, *p* < 0.001). **(D)** Diagram of 1-on-1 social interaction tests with stressed conspecifics and pretreatment with either vehicle or OTRa. **(E)** Mean (with individual replicates, normalized as percent of 3 to 5 min long test, time spent interacting with the stressed conspecific in a 1-on-1 test (PN 30: *n* = 8, PN 50: *n* = 7). OTRa significantly reduced time interacting with the stressed PN 30 conspecific but increased time interacting with the stressed PN 50 conspecific (*F*_AGE_(1, 13) = 28.66, *p* < 0.001; *F*_AGE*DRUG_(1, 13) = 32.56, *p* < 0.001). **(F)** Mean (with individual replicates) time spent interacting with a naive conspecific after intra insular cortex OT (250pg/side) or vehicle administration in a 1-on-1 social interaction (PN 30: *n* = 15; PN 50: *n* = 15). OT caused a significant increase in social interaction with naive PN 30 juveniles but a significant decrease in interaction with naive PN 50 adults (*F*_AGE*DRUG_(1, 28) = 30.08, *p* < 0.0001). To determine if PKC inhibition would interfere with social affective behavior, rats were implanted with insular cortex injection cannula and then subjected to SAP tests with pretreatment of Gö 6983 (0.5uL/side 200nM) or vehicle (10% DMSO in water). **(G)** Mean (individual replicates) time spent exploring either a naive conspecific or a stressed conspecific in the SAP test with either PN 30 (*n* = 8) or PN 50 (*n* = 7) conspecifics after intra-insula infusion of Gö 6983 antagonist. Vehicle treated experimental adult rats spent more time interacting with the stressed PN 30 juvenile conspecifics and less time with the stressed PN 50 adult conspecifics. These trends were blocked and reversed by the PKC inhibitor (*F*_STRESS*AGE*DRUG_(1, 13) = 63.75, *p* < 0.001). **(H)** Data in (G) expressed as percent preference for interaction with the stressed conspecific (Mean with individual replicates). Gö 6983 significantly reduced preference for the stressed PN 30 while increasing time spent with the stressed PN 50 conspecific (*F*_AGE*DRUG_(1, 13) = 141.10, *p* < 0.001). \**p* < 0.05, \*\**p* < 0.01, \*\*\**p <* 0.001.

Figure 5B shows the time spent interacting with the naive and stressed conspecifics and Figure 5C shows the time spent interacting with the stressed conspecifics as a percent of total interactions. In the vehicle condition, 7 of 10 experimental rats exhibited a preference for stressed over unstressed juveniles and 7 of the 8 experimental rats exhibited a preference for unstressed over stressed adults. Rats that did not exhibit preference in the vehicle condition were analyzed separately (Supplementary Fig. 5). Intra-insular cortex OTRa administration reversed these patterns resulting in a significant Age by Drug by Affect interaction (*p* < 0.001) with significantly greater time spent investigating the stressed PN 30 conspecific (vs. the naive PN30 conspecific) and less time investigating the stressed PN 50 conspecific (vs. the naïve PN50 conspecific) under vehicle conditions (*ps* = 0.001). Application of OTRa completely prevented this behavior, with more time interacting with the naive PN 30 conspecific (*p* = 0.06, approaching significance in the opposite direction of vehicle behavior) and less time with the naive PN 50 conspecific (*p* = 0.014). Percent preference scores revealed an Age by Drug interaction (*p <* 0.001) with opposing effects of OTRa in experimental rats’ behavior towards PN 30 (*p* = 0.011) and PN 50 (*p* = 0.002) groups, respectively. Thus, blockade of insular cortex OTRs prevented the age-dependent approach and avoidance behavior in response to stressed conspecifics in the SAP test.

We next sought to test the generality of the effect of OTRa using the one-on-one version of the social affective behavior test (Fig. 5D). Bilateral microinjection cannulae were placed in the insular cortex in 16 experimental rats. Rats were randomly assigned to interact with either a PN 30 juvenile or a PN 50 adult conspecific (between-subjects) and received vehicle and OTRa injections (within-subjects) in a 2 by 2 design; drug order was counter balanced by day as above. All rats received 2 social exploration tests with an unfamiliar, stressed conspecific preceded 15 min by either vehicle or OTRa microinjections. One experimental rat interacting with adult conspecifics had a misplaced cannula and was excluded. OTRa microinjection reduced the time spent interacting with the stressed PN 30 conspecifics and increased the time interacting with stressed PN 50 conspecifics in the one-on-one tests (Fig. 5E). ANOVA revealed a main effect of Age and Age by Drug interaction (*p*s < 0.001). The main effect of age indicates more interaction time with the stressed juveniles than the stressed adults, which replicates the pattern to approach stressed juveniles but avoid stressed adults that was apparent in the SAP test (as in Fig. 1H). OTRa reduced time spent interacting with the stressed juvenile over the naive juvenile (*p* < 0.001) but increased interaction with the stressed adult conspecifics over the naïve adult conspecific (*p* = 0.009). In a control experiment, OTRa had no effect on interaction between experimentally naive adult and PN 30 rats (Sup. Fig. 6).

#### Insular cortex oxytocin administration is sufficient to recapitulate social affective behaviors toward naive conspecifics

Together, the OTRa experiments demonstrate that modulation of the insular cortex by OTRs is necessary for the age-dependent approach and avoidance responses to stressed conspecifics. We next determined whether intrainsular OT administration was sufficient to increase experimental rat interaction with non-stressed conspecifics. Bilateral insular cannula implants were made in 32 rats and after recovery each received 2 one-on-one social interaction tests (3 min duration) with unfamiliar naive PN 30 or PN 50 conspecifics. A within-subjects design was used such that each rat received vehicle injections prior to one test and OT (250 pg/side; equivalent to 500nM) before the other. Injections were made 15 min before testing and drug/vehicle order was counterbalanced. One rat from each age group did not receive injections and was removed from analysis. OT microinjection increased time spent interacting with the naive PN 30 conspecific (*p* < 0.001) and decreased time with the naive PN 50 conspecific (*p* = 0.002, Fig. 5F). These findings suggest that the effect of OT in insular cortex is sufficient to bi-directionally modulate social behaviors depending upon the age of the target conspecific.

#### Social responses to stressed conspecifics require insular cortex protein kinase C (PKC)

The foregoing suggested the possibility that behavioral responses to stressed conspecifics depends upon modulation of intrinsic excitably in the insular cortex by OT and so we predicted that interference with insular cortex PKC would mimic the effect of OTRa. Rats received bilateral insular cortex cannula implants and underwent SAP tests as above with pretreatment with either vehicle or Gö 6983. Eight of 10 rats exhibited preference for the stressed PN 30 conspecific after vehicle injection and 7 of 11 rats avoided the stressed PN 50 conspecific. In these rats, Gö 6983 reversed the pattern observed under vehicle (Age by Stress by Drug interaction, *p* < 0.001). In tests with PN 30 conspecifics, experimental rats spent more time interacting with the stressed than naive conspecifics after vehicle injection (*p* < 0.001) and less time interacting with the stressed than naive conspecific after Gö injection (*p* = 0.009). In tests with PN 50 conspecifics, the experimental rats spent less time interacting with the stressed conspecific in the vehicle condition (*p* = 0.006). To summarize the behavioral pharmacology studies, experimental adult rats preferred to interact with stressed juveniles but avoided interaction with stressed adult conspecifics. In the insular cortex, blockade of the OTR and inhibition of PKC interfered with experimental rat responses to stressed conspecifics and OT administration was sufficient to reproduce the phenomena with naive conspecifics.

#### Insular cortex and the social decision-making network

As noted, the insular cortex has extensive afferent and efferent connectivity to many nodes within the SDMN, prefrontal cortex and amygdala. We therefore assessed whether the insular cortex is functionally related to these networks by using Fos immunoreactivity in 29 ROIs (for complete list and representative images see Sup. Fig. 8-9) in the 44 rats that were used for USV analysis (Fig. 1J-K) and insular Fos counts (Fig. 2B-C). Using the Brain Connectivity Toolbox ^72^, we conducted a graph theoretical analysis upon the Fos counts pooled across all treatment conditions to maximize network variation and to characterize the relationships among the ROIs. ROIs served as nodes and the rank-order correlations of Fos levels served as edges. A data-driven community detection analysis revealed that there were 2 modules within the network (Fig. 6A) in which ROIs were more highly correlated within each module than between modules (maximized modularity quotient = 0.59). The degree to which ROI Fos levels were correlated within and between modules varied across the groups, with stronger between module correlations within the stressed juvenile condition (Fig. 6B, Sup. Fig. 11). The 2 modules contain ROIs that may be categorized generally as “social” (green module) and “emotional” areas (purple module), respectively (Fig. 6C). Interestingly, insular cortex regions were found in both modules. To test whether insular cortex regions might serve as a “hub” for facilitating communication between the modules, we evaluated the participation coefficient for each node (Fig. 6D, Sup. Fig. 11); this is a measure of the degree to which nodes correlate with multiple modules and has been argued to be an appropriate measure of “hubness” for correlation-based networks ^73^. Agranular insular (AI) cortex was among the nodes with the highest participation coefficients along with prelimbic prefrontal cortex (PL), piriform cortex (Pir) and dorsal bed nucleus (BSTd), regions that are associated with emotion contagion^52^, social learning^74^ and social decision-making^6^, respectively. Thus, data-driven network analyses revealed functional integration among regions in the social brain, and that the insula is positioned to interact with previously identified nodes important for social and affective behaviors.

**Figure 6.**
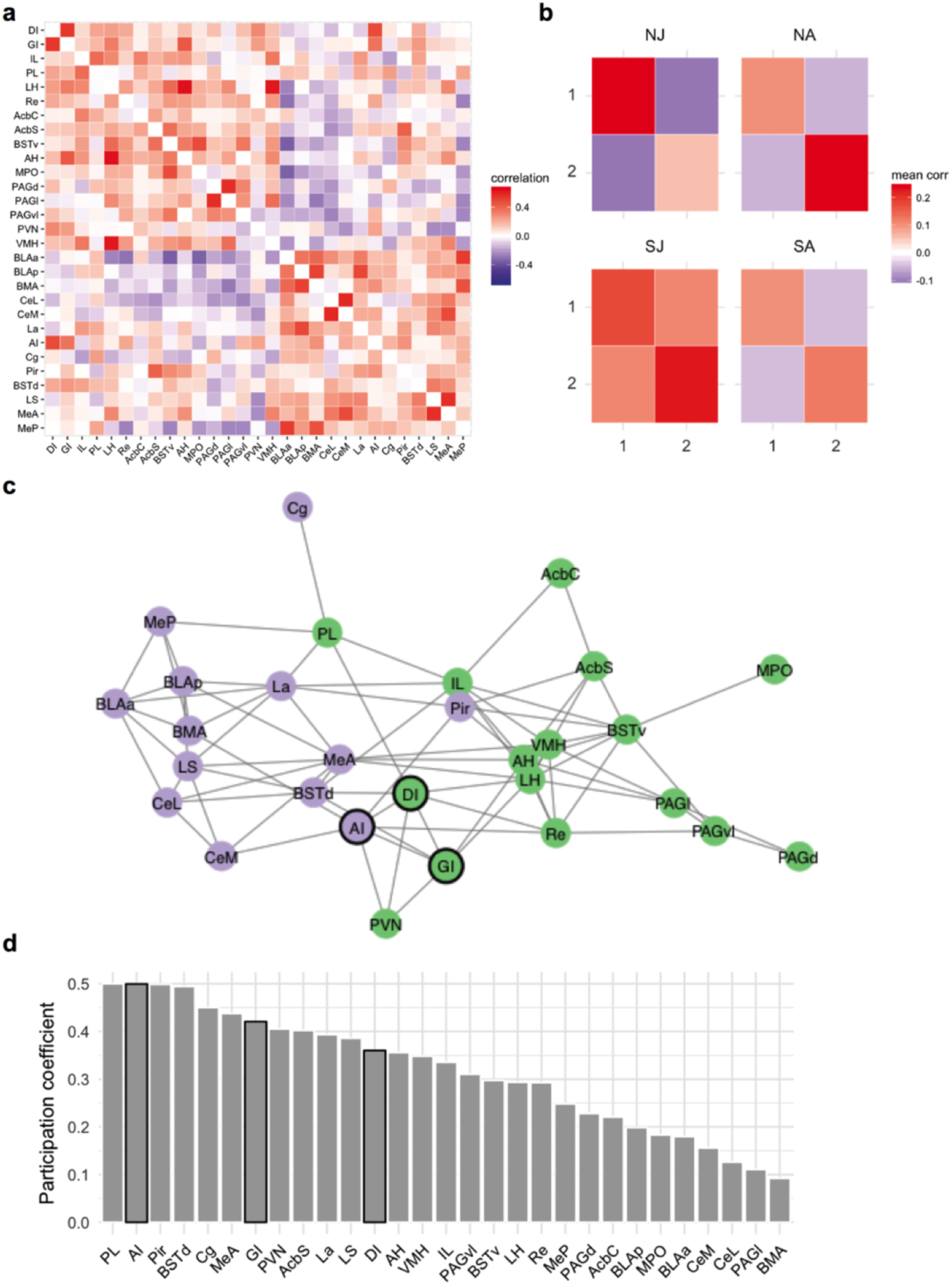
Insular Cortex and the Social Decision-Making Network. Fos immunoreactivity was determined in 29 ROIs of the rats (N=44) used in the USV quantification experiment (Fig. 1). ROIs, including the insular cortex, were selected from the SDMN and regions reported to be involved in social behavior (List of abbreviations and representative images in Sup. Figs. 8 - 9). **(A)** Correlation matrix indicating functional correlations (Kendall’s tau) among ROIs. **(B)** Graph theory based community detection analyses identified two modules. Functional correlations within and between modules 1 and 2 were averaged across ROIs and shown for each group. **(C)** Network visualization of the nodes in modules 1 (green) and 2 (purple). Here an arbitrary threshold of 0.2 was applied to facilitate network visualization, such that only edges exceeding the threshold are shown, but note that all network analyses were based on the unthresholded network. Insular cortex nodes are outlined in black. **(D)** The degree to which each node is connected to multiple functional modules was estimated by computing the participation coefficient. Bars corresponding to insular cortex ROIs are outlined in black. Network analyses by treatment group are provided in Sup. Fig. 11. NJ = naive juvenile, NA = naive adult, SJ = stressed juvenile, SA = stressed adult.

## DISCUSSION

To better understand the neural basis of social affective behavior, we developed a social affective preference (SAP) test in which the choice to approach or avoid a stressed conspecific could be quantified. In the SAP test, behaviors depended upon both the age and emotional state of the conspecific targets which may reflect a species-specific adaptation in which social stress signals are perceived and responded to as danger cues when generated by an adult, but as prosocial cues when generated by juveniles. *In vivo* pharmacological experiments, *in vitro* studies and network analyses suggest that exposure to a stressed conspecific evokes OT-dependent modulation of insular cortex output neuron excitability which orchestrates species-specific, age-dependent approach or avoidance behaviors via modulation of the social decision-making network (SDMN). The findings provide new insight into the neural basis of elementary socioemotional processes and, consistent with human neuroimaging work, warrant consideration of the insular cortex as an important component of the social brain.

A central ideal in social neuroscience is that social behaviors involve the integration of multisensory cues with situational and somatic factors ^75^. Across mammals, social communication of emotion occurs via chemosignals^76,77^, vocalizations ^65^, and overt behaviors ^78,79^. Foot-shocked rats emit social odors that can either attract conspecifics to promote social buffering or serve as social alarm signals ^80^. Although we did not investigate chemosignals, these likely contribute to the behavior of the experimental rats in the SAP test. Regarding vocalizations, 22kHz USVs are emitted as alarm signals, and frequency modulating USVs are thought to convey positive affect ^81^. Accordingly, 22kHz vocalizations were absent during naive-naive interactions, but increased dramatically when one of the conspecifics was a stressed adult. Frequency modulating (rising and trill) calls preceded bouts of social interaction (Sup. Fig. 10) and were abundant in naive-naive interactions but dropped considerably when one of the conspecifics had received footshock, a pattern consistent with a negative affective state in the stressed conspecifics. However, fewer frequency modulating calls is incongruent with the observed greater approach of experimental adults toward stressed juveniles. Perhaps stressed juveniles emit a qualitatively different type of USVs akin to the “40kHz isolation calls” which are observed in rat pups to attract the mother ^82^; this possibility will require further investigation. Regarding overt behaviors, stress increased the amount of time juvenile conspecifics engaged in self-grooming. Thus, the experimental rat may draw from a constellation of cues, including vocalizations, overt behaviors and chemical signals, to compute the age and emotional state of the conspecific. Suitably, correlations between the insular cortex and the piriform cortex and medial amygdala Fos immunoreactivtity, key olfactory processing areas modulated by OT during social interactions ^83^, were strongest after interactions with stressed juveniles (Sup. Fig. 11). Future studies will be needed to determine precisely the stimuli emitted by stressed conspecifics to determine social interactions and how this information converges on the insular cortex.

The decision to avoid a stressed adult may be adaptive as the stressed conspecific may be perceived as a social danger cue. On the other hand, it is difficult to pinpoint the motivation to approach the stressed juvenile. In human social behavior, features of the target stimulus, including age, are critical determinants to whether or not someone will approach, help or avoid another in distress ^63^ and there are examples of prosocial responses to stressed conspecifics in vole ^41^ and mouse ^59^. These prosocial examples appear to have boundary conditions. First, prosocial effects are evident in pair-bonded voles and familiar female mice, but not between strangers or in male mice; this may also generalize to rat ^84^ but see ^61^. An effect of the conspecific’s age has not previously been investigated, and our results suggest that prosocial behaviors may occur towards unfamiliar conspecifics if they are juveniles, regardless of sex (Sup. Fig. 2). Regardless of the motivation underlying the behavior of experimental rats in response to stressed conspecifics, the phenomena reported here is direct evidence of what de Waal argued to be the most elementary component of empathy: “that one party is affected by another’s emotional or arousal state (p. 282).” ^85^ The SAP test may provide a useful addition to the preclinical toolkit for investigating psychiatric diseases such as autism and schizophrenia.

Anatomically, the insular cortex is positioned to integrate social affective stimuli as a nexus between multimodal sensory inputs and emotional, executive, and social circuits ^86^. Graph theoretical analysis of Fos immunoreactivity among nodes including these three cognitive control systems revealed several features of the network underlying social affective behavior. Highly correlated patterns of activity emerged in 2 modules (Fig. 6) which were composed primarily of structures found in the SDMN such as the PAG, VMH, BST, AcbS, and PVN, structures associated with emotion, such as the nuclei of the extended amygdala. The insular cortex subfields were found in both of these modules with a high degree of participation in both networks suggesting that it is a hub between social and emotional systems. These features may be key to understanding why inhibition of the insular cortex not only interfered with social affective preferences, but in fact reversed them (Sup. Fig. 7). We hypothesize that exposure to stressed conspecifics elicits parallel activity within these modules which compete for control to either approach or avoid. When the insula is online, OT in concert with sensory cues that identify the age of the conspecific may bias the output of insula efferents to the nucleus accumbens or prefrontal cortex to promote interaction with stressed juveniles, or to the amygdala to promote avoidance in response to stressed adults as these are thought to be the proximal mediators of the rewarding, empathic, and emotional aspects of social behavior, respectively ^87^. Consonant with this view, insular OTR binding density correlated with reproductive success in male prairie voles ^88^; a behavioral endpoint that depends upon integration of social cues such as a partner’s sexual status and situational factors with the specific behaviors under the control of the SDMN.

OT altered several intrinsic properties of insular neurons that are consistent with a modulatory role, namely, reduction in AP amplitude, increase in *R*_input_, increase in input/output relationships, and reduction in sAHP; modulation of the latter 3 predict that OT could boost the output of insular cortex pyramidal neurons in response to excitatory inputs which was evident in a potentiation of evoked insular fEPSCs. Indeed, OT has been shown to increase spike output ^89–91^ and facilitate synaptic long-term potentiation ^92–94^ and our data suggest an intrinsic mechanism that may underlie shifts in synaptic efficacy and excitatory/inhibitory balance found in brain regions where OT is critical for myriad social behaviors ^67,74,90,95^. Intrinsic properties are shaped by ion channels. In preliminary whole cell voltage clamp experiments, OT reduced voltage-gated Na^+^ and L-type voltage-gated Ca^2+^ currents (data not shown) which likely mediate the reduction in AP amplitude but future studies are required to resolve the mechanism underlying the shifts in *R*_input_ and sAHP that most-likely increase in firing frequency.

Like social behavior itself, the effects of OT administration on social decision-making are sensitive to situational and interpersonal factors with OT sometimes producing prosocial effects, and at other times anti-social effects ^39^ which were both evident after OT administration to the insular cortex in the SAP test. Abnormalities in emotion recognition are central to numerous psychiatric conditions including ASD ^96–100^ and deficits in OT and insular cortex activity and connectivity are well documented correlates of symptom severity for many of these conditions ^46,101,102^. Our results suggest that disruption of insula function may be key to pathophysiology in behaviors that depend upon social decisions regarding the emotions of others. The SAP test and the network model of insular input to the SDMN presented here provide a platform for further research into the neuroanatomical and physiological systems that integrate social information with decision-making and behavior.

## ONLINE METHODS

### Rats

Male Sprague-Dawley rats were obtained from Charles River Laboratories (Wilmington, MA). Rats were allowed a minimum of 7 days to acclimate to the vivarium after arrival and housed in groups of 2-3 with free access to food and water on a 12 h light/dark cycle. Behavioral procedures were conducted within the first 4 h of the light phase. All reagents and chemicals were purchased from Fisher Scientific, Tocris or Sigma unless otherwise noted. All procedures were conducted in accordance with the Public Health Service *Guide for the Care and Use of Laboratory Animals* and were approved by the Boston College Institutional Animal Care and Use Committee.

### Social Affective Preference (SAP) Test

The SAP test allowed quantification of social interactions initiated by an adult test rat when presented simultaneously with two unfamiliar conspecific stimuli. To begin the test, the adult test subject was placed into a clear plastic cage (50 × 40 × 20 cm, L = W = H) with a wire lid. Pairs of stimuli rats were either juvenile (PN 30 +/− 2 days old) or adult (PN 50 +/− 2 days old) and were placed inside one of two clear acrylic plastic enclosures (18 × 21 = 10 cm, L = W = H) on either end of the arena. Interaction between the experimental and stimuli rats was permitted on one side of the enclosure, which consisted of clear acrylic rods, spaced 1 cm center-to-center (see photo, Fig. 1) and as in ^103^. To habituate subjects to the procedure on days 1 and 2 the adult was placed in the arena for 60 min and then empty enclosures (Day 1) or enclosures containing experimentally naive, unfamiliar stimuli (Day 2) were added for 5 min. To assess social affective preference, on Day 3 two unfamiliar stimuli were added, but one of the stimuli rats was exposed to 2 footshocks (1mA, 5 sec duration, 60 sec inter-shock-interval, Precision Regulated Animal Shocker, Coulbourn Instruments, Whitehall, PA) immediately prior to the 5 min test to induce a stressed affective state. Shock occurred in a separate room and shock parameters were selected because they were sufficient to produce a conditioned fear in our laboratory (data not shown). The 5 min test length was selected after pilot studies in which we observed a reliable decrease in social behavior after the first 5 min of test. In experiments involving optogenetics or intracerebral injections, a within-subjects design was employed such that each adult test subject was exposed to both vehicle and experimental treatments in SAP tests on consecutive days. A trained observer quantified the time spent in social interaction with each of the stimuli. Social interaction consisted of nose-nose and nose-body sniffing, and reaching into the enclosure to contact the stimulus rat. Digital video recordings were made of each test for later determination of inter-rater-reliability by a second observer completely blind to the experimental manipulations and hypotheses. Across the experiments included in this report we observed very high inter-rater reliability, *r*(80) = 0.966, *r*^*2*^ = 0.93, p < 0.0001. Although conceived independently, this paradigm has a number of features in common with the method recently reported with voles ^104^. For some tests, we also quantified a number of behaviors in the experimental adult rat from videos including time spent 1) sniffing or investigating the test arena, 2) immobile, 3) digging in the bedding, 4) self-grooming, and 5) biting or pulling at the conspecific. The first three measures were assessed to quantify general locomotor activity, self-grooming has been argued to reflect emotion contagion ^41^ and biting may reflect aggressive behaviors.

### One-on-One Social Exploration Tests

As in ^105^ each experimental subject was placed into a plastic cage with shaved wood bedding and a wire lid 60 min before the test. To begin the test a juvenile or adult was introduced to the cage for 5 min and exploratory behaviors (sniffing, pinning, and allogrooming) initiated by the adult test subject were timed by an observer blind to treatment. Juvenile and adult stimuli rats were used for multiple tests but were never used more than once for the same adult test rat. Each experimental adult was given tests on consecutive days once with an unfamiliar naive conspecific and once with an unfamiliar stressed conspecific (2 foot shocks, exactly as above); test order was counterbalanced.

### Insular Cortex Cannula Placement and Microinjection

Under inhaled isoflurane anesthesia (2-5% v/v in O_2_), cannula (26g, Plastics One, Roanoke, VA) were inserted bilaterally into the insular cortex (from Bregma: AP: −1.8mm, ML: +/−6.5mm, DV: −6.2mm from skull surface) and fixed in place with acrylic cement and stainless steel screws. Rat were administered the analgesic meloxicam (1mg/kg, Eloxiject, Henry Schein) and antibiotic penicillin G (12,000 Units, Combi-pen 48, Henry Schein) after surgery and allowed between 7-10 days recovery prior to experimentation. The OTR antagonist (OTRa) desGly-NH_2_-d(CH_2_)_5_[Tyr(Me)^2^, Thr^4^OVT ^106^ and OT were dissolved in sterile 0.9% saline vehicle, the pan-PKC inhibitor Gö 6983 was first dissolved in 100% DMSO and then diluted to 200nM in a vehicle of 10% DMSO and water. All injections were 0.5μL per side and infused at a rate of 1μL/min with an additional minute for diffusion. At the conclusion of the experiment, rats were overdosed with tribromoethanol (Sigma) and brains were dissected and sectioned at 40 µm to verify the microinjector tip location using cresyl violet stain and comparison to stereotaxic atlas ^107^. Rats with occluded injectors or having cannula located outside of the insular cortex were excluded from all analyses (See Supplementary Fig. 5).

### Optogenetics

Adult male rats underwent stereotaxic surgery to be implanted with bilateral guide cannula designed to fit a 200um optical fiber (Plastics One). After the cannula was secured, 250nL of a viral vector containing the neuronal silencing halorhodopsin eNpHr3.0 under the CamKII⍺ promoter (AAV5-CamKII⍺-eNpHR3.0-mcherry; Tye et al., 2011) or a sham virus (AAV5-CamKII⍺-YFP) was microinjected at a depth 1mm below the termination of the guide cannula at a rate of 50nL/min and allowed 5 min for diffusion. During testing, a multimodal fiber optic wire (200um core, 0.39NA, Model FT200EMT, Thorlabs) extending 1mm below the cannula tip was affixed to the stylet via a screw top ferrule (Plastics One) and connected to a laser (GL523T3-100, Shanghai Laser & Optics Century). Throughout the length of a social test green light (λ **−** 523nm) at a power of ~10-15mW/mm^2^ was administered to maintain insular inhibition for the light ON conditions. During the light OFF condition, rats underwent the social test while connected to the laser but no light was administered. Functional photo inhibition was verified in whole cell recordings of mCherry positive insular cortex pyramidal neurons in acute brain slices before, during, after green light administration (10mW/mm2) delivered through the objective of the electrophysiology microscope. The extent of transfections was determined by imaging mCherry expression with widefield fluorescent microscopy (Zeiss AxioImager Z2). Locations of transfections are provided in Supplementary Figures 5 and 6.

### Electrophysiology Solutions and Drugs

All chemicals were purchased from Fisher Scientific, Sigma-Aldrich or Tocris. Standard artificial cerebrospinal fluid (aCSF) and recording solutions were used ^108^. aCSF recording composition was (in mM) NaCl 125, KCl 2.5, NaHCO_3_ 25, NaH_2_PO_4_ 1.25, MgCl_2_ 1, CaCl_2_ 2 and Glucose 10; pH = 7.40; 310 mOsm; aCSF cutting solution was: Sucrose 75, NaCl 87, KCl 2.5, NaHCO_3_ 25, NaH_2_PO_4_ 1.25, MgCl_2_ 7, CaCl_2_ 0.5, Glucose 25 and Kynureinic acid 1; pH=7.40, 312 mOsm. The internal recording solution consisted of (in mM) K^+^-Gluconate: 115, KCl 20, HEPES 10, Mg-ATP 2, Na-GTP 0.3, and Na-Phosphocreatine 10. pH = 7.30; 278 mOsm with 0.1% biocytin. Kynurenic acid 1 mM and SR-95531 2 µM were always added to the recording aCSF to block synaptic transmission for intrinsic recordings.

### Insular cortex slices

Adult male rats were anesthetized with isofluorane, intracardially perfused with chilled (4°C), oxygenated aCSF cutting solution and quickly decapitated. 300 µm coronal slices including the insular cortex were taken using a vibratome (VT-1000S, Leica Microsystems, Nussloch, Germany). The slices were placed in oxygenated aCSF cutting solution (95% O^2^ and 5% CO^2^) at 37°C for 30 min and then at room temperature for a minimum of 30 min before slices were used for electrophysiological recordings.

### Electrophysiology

Whole-cell current-clamp recordings were obtained at 30 ± 2°C. Patch-clamp electrodes were pulled (P-1000, Sutter Instruments, CA) from 1.5 mm outer diameter borosilicate glass (Sutter Instruments, CA) and filled with intracellular solution. Electrode resistance was 3–5 MΩ in the bath and recordings were only included if the series resistance remained less than 30 MΩ with less than 10% change from baseline throughout the experiment. Slices were visualized using a 40x (0.75 N.A.) water immersion objective under infrared differential interference contrast imaging on an upright microscope (AxioExaminer D1, Zeiss, Germany). All recordings were obtained with an Axon 700B amplifier and pClamp 10 (Molecular Devices), using appropriate bridge balance and electrode-capacitance compensation. After achieving a whole-cell configuration, baseline recordings were made in aCSF until 10 minutes of stable baseline were observed, at which point 500 nM oxytocin citrate was added to the bath. The dose of 500 nM was selected after a pilot study using a range of doses from 50nM to µ1M, the largest dose reported ^95^. Because OT has high affinity for the Vasopressin 1A receptor (V1a), experiments typically isolate effects of OT to the OTR by using synthetic OTR agonists or a cocktail of OT and V1a antagonists. These steps were not taken here because although V1a receptor mRNA has been reported throughout cortex ^109^ V1a receptor binding is not evident in adult male rat insula ^110^. Analyses were performed using custom software written for Igor Pro (Wavemetrics Inc., Lake Oswego, OR).

Active properties were quantified from single spikes by holding the neuron at −67 mV, and 2.5 ms current pulses were injected to elicit a single AP. Passive properties were measured by holding the membrane potential at −67 mV and injecting 1 s current pulses through the patch electrode. The amplitudes of the current injections were between −300 pA and +400 pA in 50 pA steps. All traces in which APs were elicited were used to generate input-output curves as the total number of APs per second plotted against the injected current. EPSCs were made in the whole cell configuration with the same internal solution and aCSF with tetrodotoxin (1uM) added to some of the recordings to isolate miniature EPSCs. Recordings were made for 10 min prior to OT (10 min) and then for 10 min after OT. EPSC frequency and amplitude were determined with the mini analysis program (Synaptosoft). After recording, the slice was fixed in 4% paraformaldehyde and biocytin was visualized using the ABC method and NovaRed (Vector labs, Burlingame, CA). Only neurons with a pyramidal morphology and soma in deep layers of insular cortex were included for analysis.

Evoked field excitatory postsynaptic potentials (fEPSPs) were recorded on a 6 × 10 perforated multiple electrode array (Model: MCSMEA-S4-GR, Multichannel Systems) with integrated acquisition hardware (Model: MCSUSB60) and analyzed with MC_Rack Software (Version 3.9). Slices were placed on the array and adhered by suction of the perfusion through the perforated substrate. Bath solutions were as above and perfused through the slice from above. A stimulating electrode was selected in the deep layers of insular cortex, and fEPSPs were recorded after stimulation (0 to 5V, biphasic 220us, 500mV increments) before, during application of 500nM OT, and after (Wash). Each step in the I/O curve was repeated 3 times (20s inter-stimulus-interval) and each family of steps was replicated 3 times in each phase of the experiment. fEPSPs from channels displaying clear synaptic responses (as in Fig. 4B) and in the vicinity of the stimulating electrode were normalized to the individual channel’s maximum response to 5V stimulation at baseline; channels from the same slice were averaged for group analysis.

The electrophysiology experiments were replicated to test the dependence of OT effects on PKC. A few key aspects of the experiments were different. First, a between-groups design was used such that baseline measures were taken from all neurons after a stable recording was achieved. Then, drugs were bath applied in aCSF that contained 0.5% DMSO to permit solubility of Gö 6983 (200nM). The conditions were as follows: aCSF, aCSF with Gö 6983, OT (500nM for whole cell recordings, 1µM for fEPSPs), or OT with Gö 6983. The OT dose was increased to 1µM in the fEPSP experiment as 500nM did not reliably replicate the increase in fEPSPs when DMSO was added to the aCSF. All dependent measures were normalized to pre-drug baselines for analysis.

### Ultrasonic Vocalization Recordings and Analysis

44 adult male rats were habituated for 1h to the test arena and 24h later randomly assigned to one of 4 treatment conditions: Naive-Juvenile, Naive-Adult, Stressed-Juvenile or Stressed-Adult in a 2 by 2 (Stress by Age) design (*n* = 11/group). Rats were given 5 min, one on one social interaction tests as above with conspecifics according to their treatment group. To record USVs, an acrylic lid was placed over the arena with a 192kHz USB microphone (Ultramic192K, www.Dodotronic.com) placed directly in the center. Recordings were made using Audacity 2.1 (www.audacityteam.org), exported as .wav files and audio spectrograms were generated in Raven Pro 1.5 (The Cornell Lab of Orn-ithology https://store.birds.cornell.edu/Raven_Pro_p/ravenpro.htm). USVs were identified using the Band Limited Energy Detector function within Raven Pro. High frequency “Trill” calls were found in the range of 8-20ms, from 55-80kHz; frequency modulating “Rising” calls were 30-100ms, from 35-68kHz; and 22kHz “flat” calls were greater than 100ms, from 18-28kHz. These ranges were drawn from the literature^65^ and detection parameters were refined for each recording based on visual inspection by a trained observer who was blind to treatment condition.

### Fos analysis

After testing for USVs, each rat was left in the test cage alone and placed in a quiet room for 90 min at which point the rat was overdosed with tribromoethanol and perfused with 0.01M ice-cold heparinized phosphate buffered saline (PBS) followed by 4% paraformaldehyde. Brains were dissected and post-fixed in 4% paraformaldehyde at 4°C for 24h and transferred to 30% sucrose for 2 days. 40μm coronal slices of the entire rostral-caudal extent of the brain were collected via a cryostat at −19°C and stored in 24-well plates containing cryoprotectant at 4°C. To visualize Fos, floating sections quenched for endogenous peroxidase with 3% H_2_0_2_, blocked with 2% normal donkey serum in PBS-T (0.01% Triton-X100), and then incubated overnight in rabbit anti-c-fos antibody (1:500, Santa Cruz). Sections were then washed and incubated in biotinylated donkey anti-rabbit secondary antibody (1:200, Jackson) using the avidin-biotin complex method (ABC Elite Kit, Vector Labs) with chromogen precipitate (NovaRed Kit, Vector Laboratories). Sections were floated onto glass slides, dehydrated, cleared, and coverslipped with Permount. All steps were conducted at room temperature. To quantify Fos immunoreactive nuclei, tissue was imaged on a Zeiss Axioimager Z2 light microscope. Tiled images containing the ROIs were taken using a Zeiss AxioCam HRc digital camera through a 10x objective (N.A. 0.45). Representative images are provided in Supplementary Figure 8. Using ImageJ software, images were converted to 16-bit, ROIs were traced with reference to the rat brain atlas, and Fos immunoreactivity was quantified using the cell counter plug-in using parameters that were validated by comparison to manual counts by a trained observer. Cell density was computed the number of Fos immunoreactive cells divided by the ROI area (in pixels) for ANOVA and network analyses.

### Statistical Analysis

Sample sizes were initially determined based on prior work using social interaction ^105,111^ and intrinsic physiology ^68^. For all of the experiments that entailed a mechanistic manipulation of the insular cortex including optogenetics, OT, OTRa, and PKC inhibitor infusions, we observed a portion of rats that did not exhibit the expected behavior towards stress conspecifics. Therefore, rats were excluded from the statistical analysis if any of the following conditions were met: 1) they did not express the expected preference for stressed juveniles or avoidance of stressed adults was greater or less than 50% in the control condition, 2) the cannula were occluded or found to be outside of the insula, and/or 3) virus expression was found to be unilateral or outside of the insula. The data from rats excluded due item 1 above are provided for inspection in Supplementary Figure 5. To compare differences between mean scores of social interaction and electrophysiological endpoints we used ttests and analysis of variance (ANOVA). Individual replicate data are provided in the figures. In most experiments, there were within-subjects variables, which were treated as such in the analysis (paired samples t-test or repeated measures ANOVA). Main effects and interactions were deemed significant when *p* < 0.05 and all reported post hoc test *p* values are Sidak-adjusted, to maintain an experiment-wise risk of type I errors at α = 0.05. ANOVA analyses were conducted in Prism 7.0c (GraphPad Software) and SPSS Statistics 24 (IBM).

Graph theoretical analysis has been applied to Fos datasets to characterize functional relationships in rodent neural circuits ^112–115^. Network analyses were conducted with the Rubinov & Sporns (2010) Brain Connectivity Toolbox (https://sites.google.com/site/bctnet/) in MATLAB R2015a (The Mathworks Inc.). Kendall’s rank correlation was used to evaluate the relationships between each pair of the 29 included ROIs (nodes), computed across the sample of 44 rats that were used for USV analysis. Community detection analyses used a spectral community detection algorithm ^116^ applied to the weighted, unthresholded correlation network. Participation coefficients were calculated to determine the degree to which nodes participated in multiple networks, based on arguments that this measure is the most appropriate way to identify “hubs” in correlation-based networks ^73^. Participation coefficients were based on positive edges only. Network visualization was conducted in R version 3.2.4 with the ggplot2 ^117^ and ggnet ^118^ packages. Network analysis scripts are freely available at https://github.com/memobc and complete datasets are available by contacting the authors.

## Author Contributions

Conceptualization, J.P.C., J.A.V., M.M.R-C., M.R.; Methodology, M.M.R-C., J.A.V., M.R. J.P.C.; Investigation; J.A.V., M.M.R-C., K.B.G., A.P., M.M., M.R., J.P.C.; Writing – Original Draft, J.A.V., M.M.R-C., and J.P.C.; Writing -- Revision & Editing, M.M.R-C., M.R., J.P.C.; Funding Acquisition, J.P.C.; M.M.R-C.

## Acknowledgements

The authors wish to thank Dr. Maurice Manning for kindly providing OTRa, Dr. Karl Deisseroth for making optogenetic vectors freely available, and Drs. Alexa Veenema and Daniel Adams for constructive discussions related to the project. Funding for this work was provided by the Boston College Undergraduate Research Fellowship, National Science Foundation Grant #1258923, NIH Grant MH093412 and the Brain and Behavior Research Foundation grant No. 19417.

## FIGURE LEGENDS

**Supplementary Video 1.** Social Affective Preference Test. In this example, the experimental adult was presented with a stressed PN 30 conspecific in the left chamber and a naive PN 30 conspecific on the right. The appearance of a green rectangle identifies bouts of social interaction with the stressed conspecific.

